# 17α-ethynylestradiol (EE2) limits the impact of ibuprofen upon respiration by streambed biofilms in a sub-urban stream

**DOI:** 10.1101/718924

**Authors:** Peter McClean, William Ross Hunter

## Abstract

Pharmaceutical compounds such as the non-steroidal anti-inflammatory drug ibuprofen and the artificial estrogen 17α-ethynylestradiol (EE2) are contaminants of emerging concern in freshwater systems. Globally, human pharmaceutical use is growing by around ∼3 % per year, yet we know little about how interactions between different pharmaceuticals may affect aquatic ecosystems. Here we test how interactions between ibuprofen and EE2 affect the growth and respiration of streambed biofilms. We used contaminant exposure experiments to quantify how these compounds affected biofilm growth (biomass), respiration, net primary production (NPP) and gross primary production (GPP), both individually and in combination. We found no effects of either ibuprofen or EE2 on biofilm biomass (using ash free dry mass as a proxy) or gross primary production. Ibuprofen significantly reduced biofilm respiration and altered NPP. Concomitant exposure to EE2, however, counteracted the inhibitory effects of ibuprofen upon biofilm respiration. Our study, thus, demonstrates that interactions between pharmaceuticals in the environment may have complex effects upon microbial contributions to aquatic ecosystem functioning.

## 1. Introduction

Human pharmaceuticals and personal care products (PPCPS) are contaminants of emerging concern within the environment (Rosi-Marshall and Royer 2012; Gaston et al. 2019). Since the year 2000, pharmaceutical use has grown by approximately 3% per year globally and this is predicted to increase further as human populations grow (Van Broeckel et al. 2014). Removal of PPCPs via waste-water treatment plants (WWTPs) is inefficient leading to the constant release of low doses of compounds such as non-steroidal anti-inflammatory drugs (NSAIDs) (e.g. ibuprofen), antimicrobial compounds (e.g. triclosan, and trimethoprim) and artificial estrogens (e.g. 17α-ethynylestradiol) into the aquatic environment (Gros et al. 2007; Álvarez-Muñoz et al. 2015; Archer et al. 2017). This is potentially problematic because these compounds are specifically designed specifically to produce physiological effects within an organism, at ultra-low (nano-molar) concentrations (Rosi-Marshall and Royer, 2012; Van Broeckel et al. 2014; Álvarez-Muñoz et al. 2015). Eco-toxicological studies reveal that PPCPs at environmental concentrations can have significant physiological effects on both aquatic fauna and microorganisms, with the potential to disrupt aquatic ecosystem functioning altering carbon and nutrient cycling, and negatively affect water quality (Jobling et al. 2003; Hernando et al. 2006; Rosi-Marshall et al. 2013; Drury et al. 2013; Źur et al. 2018; Gallagher and Reisinger, 2020).

In headwater streams, aquatic biofilms attached to the streambed represent the dominant mode of microbial life Besemer et al. 2012; Battin et al. 2016). Biofilms, composed of consortia of bacteria and unicellular eukaryotic algae bound within a complex matrix of extracellular polymeric substances (EPS), play a key role in the functioning of fluvial ecosystems, controlling both the transport and degradation of organic matter within a stream (Battin et al. 2016). Rosi-Marshall et al. (2013) revealed that aquatic PPCPs such as caffeine, cimetidine, ciprofloxacin, diphenhydramine, metformin and ranitidine had negative effects upon biofilm growth, respiration, and community composition. PPCPs, however, are diverse group of chemicals, which may interact with each other in a multitude of different, and often-unexpected ways (Rosi-Marshall et al. 2013; Gerbersdorf et al. 2015; Gaston et al. 2019; Robson et al., 2020). Consequently, a mechanistic understanding of the interactions between different PPCPs is needed if we are to fully understand their environmental impacts.

Within the broad spectrum of PPCPs the non-steroidal anti-inflammatories (NSAIDs), such as ibuprofen, and artificial estrogens, such as 17α-ethynylestradiol, represent some of the most commonly detected compounds in aquatic systems (Álvarez-Muñoz et al. 2015; Gaston et al. 2019). NSAIDs are known to have antimicrobial properties, with ibuprofen exhibiting potential as a biofilm control agent (Reslinski et al. 2015; Shah et al. 2018; Źur et al. 2018; Oliveira et al. 2019). Conversely, artificial estrogens and other endocrine disruptors may adsorb onto microbial biofilms facilitating their biological degradation (Writer et al. 2012; Zhang et al. 2014; Adeel et al. 2017). There are currently no known therapeutic interactions between NSAIDs and artificial estrogens in animal systems. The fact that these compounds elicit different microbial responses, however, suggests there may be potential for interactions between NSAIDs and artificial estrogens to affect the growth and metabolism of aquatic microorganisms. Here we present the first data on how interactions between ibuprofen and 17α-ethynylestradiol (hereafter, EE2) affect the growth and respiration of streambed biofilms. We *conducted in situ* contaminant exposure experiments, following Costello et al. (2016), to test how chronic exposure to ibuprofen, and EE2, both individually and in combination, affected streambed biofilm growth, primary production and respiration.

## 2. Materials and Methods

All experiments were carried out between the 30^th^ November 2018 and the 22^nd^ January 2019 in the Ballysally Blagh (Latitude: 55°08’45.1"N Longitude: 6°40’18.0"W), a ground-water fed second-order stream. The Ballysally Blagh is a tributary of the lower River Bann (Northern Ireland), draining a mixed agricultural (consisting of 21.9 % arable; 55.9 % grassland; 13.7 % heathland; 1.9 % woodland) and urban (7.3 %) catchment of 14.2 km^2^. The mean volumetric rate for water flow in the Ballysally Blagh is 0.21 (± 0.27) m^3^ s^-1^, measured at a V-shaped weir (National River Flow Archive. 2019) and the stream is defined as eutrophic, with dissolved nitrate concentrations ranging between 1.37 and 14.15 ml.l^-1^ and soluble reactive phosphorus concentrations between 0.033 and 0.4 mg.l^-1^. Water temperature at the study site was recorded at 1-hour intervals throughout the experiment using a HOBO MX2204 Bluetooth temperature logger. Temperatures ranged between 9.35 ^°^C and 5.16 ^°^C, with a mean temperature of 7.72 (± 0.85) ^°^C recorded over the study period.

Contaminant exposure experiments were conducted following Costello et al. (2016). Briefly, forty 120 ml screw cap sample pots where filled with 2 % agar, of which ten were spiked with a fixed 0.5 mmol.l^-1^ concentration of ibuprofen, ten with a fixed 0.5 mmol.l^-1^ concentration of of EE2, ten spiked with fixed 0.5 mmol.l^-1^ concentrations of both ibuprofen and EE2, and ten received no pharmaceutical treatment (control). Both ibuprofen and EE2 have relatively low solubility in water (21 mg.l^-1^ and 3.6 mg.l^-1^ respectively). As such, stock solutions for each pharmaceutical treatment were made up by dissolving 159 mg of ibuprofen (Sigma-Aldrich, Product No. I4883), 105 mg of EE2 (Sigma-Aldrich, Product No. E4876) or both in 11 ml of 70 % ethanol. 1 ml aliquots of the stock solution were then used to dose each contaminant exposure experiment and the control treatments receiving a 1 ml aliquot of 70 % ethanol. Pre-combusted Whatman® 45 mm GF/F filters were placed onto of the solid agar and secured using the screw cap, to provide a substratum for streambed biofilm colonization. Contaminant exposure experiments were then secured to four L-shaped metal bars (*l* = 1000 mm; *w* = 50 mm; *d* = 50 mm) and deployed at 10 cm depth, in an area of turbulent flow (riffle) within the stream.

Environmental chambers were assembled from two Curry’s Essentials® C61CF13 chest freezers, with the power source re-routed through Inkbird ITC-308 Digital Temperature Controller used to override the freezers internal thermostat. A single Tetra HT50 (50 Watt) aquarium heater was also attached to the Inkbird temperature controller of each unit to help stablise the internal temperature. Two NICREW planted aquarium LED strip lights were attached to the lid, providing a source of photosynthetically active radiation (– 106.0 μmol m^-2^ s^-1^, measured using an Apogee Instruments Photosynthetically Active Radiation Meter). Environmental chambers were filled with 20 l of streamwater and the internal temperatures set to 7.7 °C. The contaminant exposure experiments were left *in situ* for 54 days, after which they were recovered from the stream, directly placed into one of the environmental chambers and allowed to acclimate over 24 hours. During the acclimation period each mesocosm was aerated using a Aqualine Hailea Aco-9630.

After the acclimation period, biofilm respiration and gross primary production were determined by changes in oxygen consumption by enclosing each contaminant exposure experiment into a sealed transparent Perspex® push core (height = 30 cm, internal diameter = 7 cm) chambers, containing 1 litre of sterile-filtered streamwater and held at 7.7 °C in one of the environmental chambers (Bott et al. 1978; Fellows et al. 2006). Biofilm respiration (R) were quantified by measuring the change in oxygen concentrations over a one-hour period (oxygen consumption in darkness (PAR ∼ 0.0 μmol m^-2^ s^-1^) using a Hach Sension 6 dissolved oxygen meter. Biofilm net primary productivity (NPP) was then quantified by measuring the change in oxygen concentration over a one 1-hour period, under artificial illumination (PAR ∼ 106.0 μmol m^-2^ s^-1^). Biofilm gross primary productivity (GPP) by the biofilms was then calculated from NPP and R as:

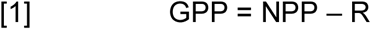

In cases where NPP was more negative that R (indicating greater oxygen consumption under artificial illumination) the baseline for GPP defaulted to zero.

Microbial biomass within each Contaminant Exposure Experiment was quantified as Ash Free Dry Mass of the GF/F filters. These were dried for 48 hours at 65 ^°^C and then subsequently combusted at 550 ^°^C for 2 hours. We estimated the daily dose of the pharmaceuticals delivered within each treatment following Costello et al. (2016), assuming that ibuprofen and EE2 doses were proportional to the agar mass lost.

All data are available in the supplementary information. Data analyses were conducted in the R statistical computing environment using the *base* and *ggplot2* packages (R Development Core Team. 2009; Wickham, 2016). We tested for independent and combined effects of ibuprofen and EE2 upon microbial biomass (Ash Free Dry Weight), Respiration and NPP and GPP using two-way analysis of variance (ANOVA). Post-hoc testing of significant interactions was conducted using Tukey’s test for Honest Significant Difference. All data were visually explored, to ensure they conformed to the assumptions of normality and homoscedacity, following Zuur et al. (2010). Microbial biomass data were log_10_ transformed to ensure the residuals of the ANOVA model conformed to a normal distribution.

## 3. Results

Using ash free dry mass as a proxy for microbial biomass we detected no significant effects (*p* > 0.05) of pharmaceutical exposure upon microbial biofilm growth (Fig 1 A; Table 1 a). We detected a significant interaction (*p* < 0.001_df = 1; F = 18.75_) between ibuprofen and EE2 affecting microbial respiration (Fig 1 B; Table 1 b). Exposure to ibuprofen alone inhibited microbial oxygen consumption by ∼ 38 %, whilst exposure to EE2 alone resulted in a slight (non-significant) increase in oxygen consumption of ∼ 5 %. In combination, EE2 counteracted the inhibitory effect of ibuprofen upon of microbial respiration, resulting in no significant change in respiration relative to the control. Biofilm NPP was negative in all treatments, with ibuprofen exposure resulting in a significant decrease in oxygen consumption (*p* = 0.009_df = 1; F = 7.483_), reflecting the effect on biofilm respiration (Fig 1 C; Table 1 c). Across all treatments GPP was close to zero, with no significant effects (*p* > 0.05) of either ibuprofen or EE2 We did, however, detect a non-significant increase in oxygen production by biofilms exposed to both ibuprofen and GPP (Fig 1 D; Table 1 d).

**Table 1.** ANOVA summary tables of the effects of Ibuprofen and 17α-ethynylestradiol [EE2] upon a) biomass (ash free dry weight), b) respiration, c) Net Primary Production and d) Gross Primary Production of cultured streambed biofilms.

| a) Biomass (Ash Free Dry Weight) |  |  |  |  |  |
| --- | --- | --- | --- | --- | --- |
|  | Df | SS | MS | F | *p* |
| Ibuprofen | 1 | 0.001 | 0.00086 | 0.008 | 0.931 |
| EE2 | 1 | 0.006 | 0.00586 | 0.051 | 0.822 |
| Ibuprofen : EE2 | 1 | 0.151 | 0.15142 | 1.331 | 0.256 |
| Residuals | 36 | 4.097 | 0.11379 |  |  |
| b) Respiration |  |  |  |  |  |
|  | Df | SS | MS | F | *p* |
| Ibuprofen | 1 | 6482 | 6482 | 41.13 | **<0.001** |
| EE2 | 1 | 5085 | 5085 | 32.26 | **<0.001** |
| Ibuprofen : EE2 | 1 | 2952 | 2952 | 18.73 | **<0.001** |
| Residuals | 36 | 5674 | 158 |  |  |
| c) Net Primary Production |  |  |  |  |  |
|  | Df | SS | MS | F | *p* |
| Ibuprofen | 1 | 7546 | 7545.8 | 7.483 | **0.009** |
| EE2 | 1 | 408 | 408.1 | 0.405 | 0.528 |
| Ibuprofen : EE2 | 1 | 38 | 38.2 | 0.038 | 0.847 |
| Residuals | 36 | 36303 | 1008.4 |  |  |
| d) Gross Primary Production |  |  |  |  |  |
|  | Df | SS | MS | F | *p* |
| Ibuprofen | 1 | 931.5 | 931.46 | 1.737 | 0.196 |
| EE2 | 1 | 1201.0 | 1200.96 | 2.240 | 0.143 |
| Ibuprofen : EE2 | 1 | 1607.4 | 1607.40 | 2.998 | 0.092 |
| Residuals | 36 b | 19300.8 | 536.13 |  |  |

**Figure 1.**
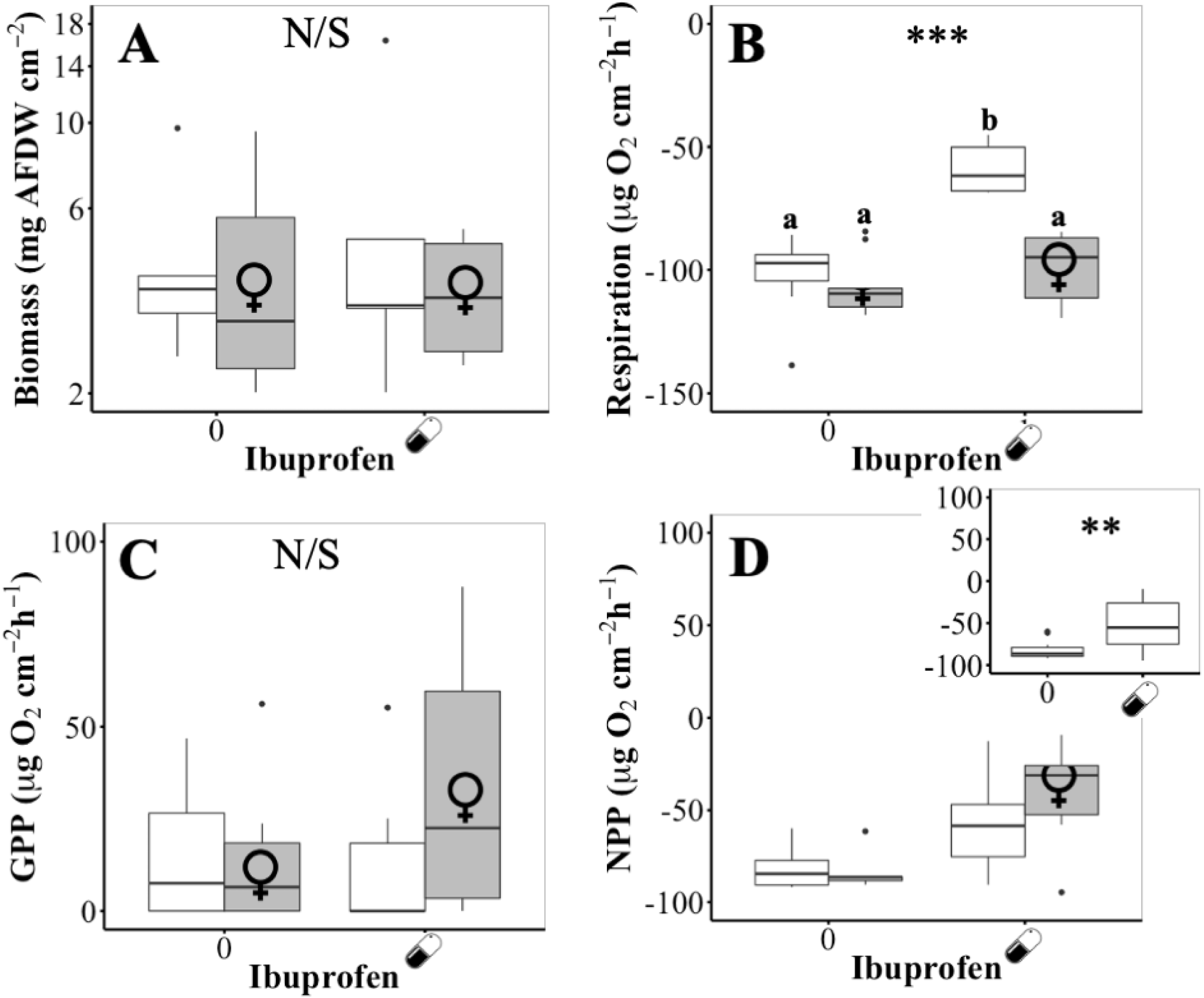
Effects of Ibuprofen () and 17α-ethynylestradiol () upon the (A) biomass (ash free dry weight), (B) respiration and (C) Net Primary Production and (D)Gross Primary Production of cultured streambed biofilms. Significance levels: \*\*\**p* < 0.001; \*\**p* < 0.01; \**p* < 0.05; N/S *p* > 0.05. Where significant interactions were identified, groups labelled with the same lowercase letter are not significantly different (*p* > 0.05; Tukey’s tests).

## 4. Discussion

Our study demonstrates that interactions between the NSAID ibuprofen and the artificial estrogen EE2 have a significant effect upon the streambed biofilm respiration. Specifically, concomitant exposure to both ibuprofen and EE2 reduced the depressive effect of ibuprofen upon biofilm respiration. Ibuprofen is known to have antimicrobial properties and has been reported to inhibit biofilm formation by both *Staphylococcus aureus* and *Escherichia coli* (Reśliński et al. 2015; Shah et al. 2018; Oliviera et al. 2019). It is, therefore, unsurprising that ibuprofen inhibited microbial respiration within the streambed biofilms. EE2 has been observed to adsorb to microbial biofilms (Writer et al. 2012) where it can then be used by the resident microorganisms as an organic matter source Stumpe et al. 2009; Ribeiro et al. 2010). Consequently, biofilms have been proposed as a tool for the removal of artificial estrogens and other endocrine disruptors within wastewater treatment facilities (Pieper and Rotard, 2011). The presence of EE2 as an energy source may, therefore, counteract the inhibitory effects of ibuprofen (Combalbert et al., 2010), whilst sorption of EE2 to the biofilm matrix may protect the microbial cells by reducing the space available onto which ibuprofen molecules may bind (Writer et al. 2012; Zhang et al. 2014). These mechanisms, however, remain speculative and require further investigation within controlled laboratory experiments.

The negative NPP within the experiment suggests that our biofilms were heterotrophic, relying on organic matter from the surrounding environment to provide energy and nutrients for biofilm growth. The significant effects of ibuprofen upon NPP, therefore, provide further evidence of this specific PPCP inhibits heterotrophic metabolism in streambed biofilms. Autotrophic activity was low, within our study, limiting our ability to infer how either ibuprofen or EE2 affects the algal component within our biofilms. Nevertheless, the non-significant increase in GPP within biofilms exposed to both pharmaceuticals further suggests that EE2 may mediates microbial responses ibuprofen exposure. This experiment was, however, conducted during the winter, when algal growth within streambed biofilms is typically low (e.g. Duncan and Blinn, 1989; Francoeur et al., 1999). To adequately test how interactions between ibuprofen and EE2 affect autrophic biofilms, requires repetition of the study during spring or summer, when longer day length is likely to promote higher algal growth at the streambed.

Given ibuprofen’s potential as a biofilm control agent (Reslinski et al. 2015; Shah et al. 2018; Źur et al. 2018; Oliveira et al. 2019), we were surprised to observe that it had no effect upon biofilm biomass within our experiments. This, however, may reflect the development of microbial resistance to anthropogenic stressors such as pharmaceuticals in agricultural and urban catchments to (e.g. Drury et al., 2013; Cai et al., 2016; Qu et al, 2017; Roberto et al. 2018). Furthermore, siltation of fine particulate matter may affect the accuracy of ash free dry mass as a measure of biomass in urban and agricultural streams. This leads us to suggest that complimentary analysis of specific microbial biomarkers, such as polar lipid fatty acids (Middelburg et al 2000; Frostegård et al., 2010; Hunter et al., 2012, 2013) and extracellular polysaccharide quantification (Fish et al., 2017; Grzegorczyk et al., 2018) may provide further insight how these pharmaceutical may affect biofilm biomass and structure.

Within this short paper we demonstrate that interactions between NSAIDs and artificial estrogens could have important implications for aquatic ecosystem functioning during the winter, when lower water temperatures limit microbial activity within streambed biofilms (Ylla et al. 2012). Whilst the doses of ibuprofen and EE2 within our study appear high, they are broadly comparable with doses used in many other contaminant exposure experiments (Drury et al., 2013; Rosi-Marshall et al. 2013, Rosi et al., 2018; Gallagher and Reisinger 2020). Our experiment, thus, provides a reasonably realistic insight into of how interactions between these two PPCPs affect aquatic microbial activity.

Our study supports a growing body of evidence suggesting that PPCPs represent a major threat to ecosystem functioning in many streams and rivers (Jobling et al. 2003; Hernando et al. 2006; Gros et al. 2007; Rosi-Marshall and Royer, 2012; Rosi-Marshall et al. 2013; Álvarez-Muñoz et al. 2015; Ruhí et al. 2016; Archer et al 2017). Interactions between PPCPs and their effects within the environment are potentially complex and mediated by changes in environmental context (Rosi-Marshall et al. 2013; Rosi et al., 2018 Gallagher and Reisinger, 2020). Future studies need to investigate how the interactions between different PPCPs affect aquatic microbial communities under different regimes of temperature, aquatic chemistry and ecological community structure. This demands the design of field-based contaminant exposure experiments that test the interactions between a range of PPCPs both within and between freshwater catchments. Here, we also highlight the need to identify what underlying biochemical mechanisms determine how interactions between different PPCPs affect aquatic microbial processes.

## 5. Acknowledgments

This work was completed by PMcC during his final year undergraduate research project, supervised by WRH. It was funded through start-up funds provided to WRH by the University of Ulster’s School of Geography and Environmental Science. We acknowledge fieldwork assistance by Ashley Williamson, and technical support in the lab from Peter Devlin and Hugo McGrogan.

## Conflicts of Interest

The authors declare no conflicts of interest relating to this study.

## Data Accessibility

All data related to this publication are available as a supplementary data file alongside this paper.

## Notes

https://pure.ulster.ac.uk/en/datasets/dataset-supporting-mcclean-and-hunter-17a-estradiol-limits-the-im

### Competing Interest Statement

The authors have declared no competing interest.

### Summary of Updates

This version of the manuscript has been revised to update the introduction, clarify the methods and the interpretation of the results within the discussion, following journal peer-review. The revised paper is now, once again, under review.

