## Supplementary Data for "17α-ethynylestradiol (EE2) limits the impact of ibuprofen upon respiration by streambed biofilms in a sub-urban stream" (from another version of this preprint: version 2, 2020-03-31)

Supplementary Table 1: Hourly temperature readings during the biofilm growth phase of the experiment.

| Date Time - UTC | Temp, (*C) |
| --- | --- |
| 30/11/2018 10:27 | 8.57 |
| 30/11/2018 11:27 | 8.65 |
| 30/11/2018 12:27 | 8.61 |
| 30/11/2018 13:27 | 8.55 |
| 30/11/2018 14:27 | 8.45 |
| 30/11/2018 15:27 | 8.42 |
| 30/11/2018 16:27 | 8.34 |
| 30/11/2018 17:27 | 8.27 |
| 30/11/2018 18:27 | 8.18 |
| 30/11/2018 19:27 | 8.11 |
| 30/11/2018 20:27 | 8.02 |
| 30/11/2018 21:27 | 7.97 |
| 30/11/2018 22:27 | 7.9 |
| 30/11/2018 23:27 | 7.83 |
| 01/12/2018 00:27 | 7.79 |
| 01/12/2018 01:27 | 7.72 |
| 01/12/2018 02:27 | 7.65 |
| 01/12/2018 03:27 | 7.57 |
| 01/12/2018 04:27 | 7.48 |
| 01/12/2018 05:27 | 7.4 |
| 01/12/2018 06:27 | 7.37 |
| 01/12/2018 07:27 | 7.39 |
| 01/12/2018 08:27 | 7.46 |
| 01/12/2018 09:27 | 7.56 |
| 01/12/2018 10:27 | 7.62 |
| 01/12/2018 11:27 | 7.63 |
| 01/12/2018 12:27 | 7.67 |
| 01/12/2018 13:27 | 7.55 |
| 01/12/2018 14:27 | 7.41 |
| 01/12/2018 15:27 | 7.29 |
| 01/12/2018 16:27 | 7.18 |
| 01/12/2018 17:27 | 7.11 |
| 01/12/2018 18:27 | 7.03 |
| 01/12/2018 19:27 | 7.01 |
| 01/12/2018 20:27 | 7.01 |
| 01/12/2018 21:27 | 6.98 |
| 01/12/2018 22:27 | 7.01 |
| 01/12/2018 23:27 | 7.03 |
| 02/12/2018 00:27 | 7.03 |
| 02/12/2018 01:27 | 7 |
| 02/12/2018 02:27 | 6.95 |
| 02/12/2018 03:27 | 6.99 |
| 02/12/2018 04:27 | 7.03 |
| 02/12/2018 05:27 | 7.08 |
| 02/12/2018 06:27 | 7.18 |
| 02/12/2018 07:27 | 7.31 |
| 02/12/2018 08:27 | 7.37 |
| 02/12/2018 09:27 | 7.54 |
| 02/12/2018 10:27 | 7.74 |
| 02/12/2018 11:27 | 7.92 |
| 02/12/2018 12:27 | 8.02 |
| 02/12/2018 13:27 | 8.11 |
| 02/12/2018 14:27 | 8.15 |
| 02/12/2018 15:27 | 8.2 |
| 02/12/2018 16:27 | 8.23 |
| 02/12/2018 17:27 | 8.28 |
| 02/12/2018 18:27 | 8.3 |
| 02/12/2018 19:27 | 8.33 |
| 02/12/2018 20:27 | 8.36 |
| 02/12/2018 21:27 | 8.37 |
| 02/12/2018 22:27 | 8.38 |
| 02/12/2018 23:27 | 8.41 |
| 03/12/2018 00:27 | 8.42 |
| 03/12/2018 01:27 | 8.43 |
| 03/12/2018 02:27 | 8.38 |
| 03/12/2018 03:27 | 8.36 |
| 03/12/2018 04:27 | 8.35 |
| 03/12/2018 05:27 | 8.32 |
| 03/12/2018 06:27 | 8.28 |
| 03/12/2018 07:27 | 8.23 |
| 03/12/2018 08:27 | 8.26 |
| 03/12/2018 09:27 | 8.33 |
| 03/12/2018 10:27 | 8.44 |
| 03/12/2018 11:27 | 8.51 |
| 03/12/2018 12:27 | 8.58 |
| 03/12/2018 13:27 | 8.59 |
| 03/12/2018 14:27 | 8.58 |
| 03/12/2018 15:27 | 8.41 |
| 03/12/2018 16:27 | 8.36 |
| 03/12/2018 17:27 | 8.39 |
| 03/12/2018 18:27 | 8.32 |
| 03/12/2018 19:27 | 8.27 |
| 03/12/2018 20:27 | 8.24 |
| 03/12/2018 21:27 | 8.19 |
| 03/12/2018 22:27 | 8.16 |
| 03/12/2018 23:27 | 8.14 |
| 04/12/2018 00:27 | 8.05 |
| 04/12/2018 01:27 | 7.96 |
| 04/12/2018 02:27 | 7.85 |
| 04/12/2018 03:27 | 7.75 |
| 04/12/2018 04:27 | 7.6 |
| 04/12/2018 05:27 | 7.48 |
| 04/12/2018 06:27 | 7.38 |
| 04/12/2018 07:27 | 7.34 |
| 04/12/2018 08:27 | 7.47 |
| 04/12/2018 09:27 | 7.63 |
| 04/12/2018 10:27 | 7.75 |
| 04/12/2018 11:27 | 7.92 |
| 04/12/2018 12:27 | 8.02 |
| 04/12/2018 13:27 | 7.98 |
| 04/12/2018 14:27 | 8 |
| 04/12/2018 15:27 | 7.94 |
| 04/12/2018 16:27 | 7.85 |
| 04/12/2018 17:27 | 7.77 |
| 04/12/2018 18:27 | 7.7 |
| 04/12/2018 19:27 | 7.58 |
| 04/12/2018 20:27 | 7.44 |
| 04/12/2018 21:27 | 7.32 |
| 04/12/2018 22:27 | 7.2 |
| 04/12/2018 23:27 | 7.09 |
| 05/12/2018 00:27 | 6.98 |
| 05/12/2018 01:27 | 6.93 |
| 05/12/2018 02:27 | 6.87 |
| 05/12/2018 03:27 | 6.81 |
| 05/12/2018 04:27 | 6.76 |
| 05/12/2018 05:27 | 6.73 |
| 05/12/2018 06:27 | 6.7 |
| 05/12/2018 07:27 | 6.7 |
| 05/12/2018 08:27 | 6.78 |
| 05/12/2018 09:27 | 6.89 |
| 05/12/2018 10:27 | 6.99 |
| 05/12/2018 11:27 | 7.05 |
| 05/12/2018 12:27 | 7.09 |
| 05/12/2018 13:27 | 7 |
| 05/12/2018 14:27 | 6.94 |
| 05/12/2018 15:27 | 6.84 |
| 05/12/2018 16:27 | 6.72 |
| 05/12/2018 17:27 | 6.66 |
| 05/12/2018 18:27 | 6.61 |
| 05/12/2018 19:27 | 6.55 |
| 05/12/2018 20:27 | 6.48 |
| 05/12/2018 21:27 | 6.39 |
| 05/12/2018 22:27 | 6.36 |
| 05/12/2018 23:27 | 6.31 |
| 06/12/2018 00:27 | 6.35 |
| 06/12/2018 01:27 | 6.48 |
| 06/12/2018 02:27 | 6.61 |
| 06/12/2018 03:27 | 6.76 |
| 06/12/2018 04:27 | 6.97 |
| 06/12/2018 05:27 | 7.16 |
| 06/12/2018 06:27 | 7.16 |
| 06/12/2018 07:27 | 7.25 |
| 06/12/2018 08:27 | 7.37 |
| 06/12/2018 09:27 | 7.46 |
| 06/12/2018 10:27 | 7.65 |
| 06/12/2018 11:27 | 7.83 |
| 06/12/2018 12:27 | 7.93 |
| 06/12/2018 13:27 | 7.99 |
| 06/12/2018 14:27 | 8.02 |
| 06/12/2018 15:27 | 8.02 |
| 06/12/2018 16:27 | 8.05 |
| 06/12/2018 17:27 | 8.03 |
| 06/12/2018 18:27 | 8.05 |
| 06/12/2018 19:27 | 8.04 |
| 06/12/2018 20:27 | 8.06 |
| 06/12/2018 21:27 | 8.06 |
| 06/12/2018 22:27 | 8.07 |
| 06/12/2018 23:27 | 8.11 |
| 07/12/2018 00:27 | 8.14 |
| 07/12/2018 01:27 | 8.16 |
| 07/12/2018 02:27 | 8.24 |
| 07/12/2018 03:27 | 8.35 |
| 07/12/2018 04:27 | 8.49 |
| 07/12/2018 05:27 | 8.56 |
| 07/12/2018 06:27 | 8.63 |
| 07/12/2018 07:27 | 8.76 |
| 07/12/2018 08:27 | 8.83 |
| 07/12/2018 09:27 | 8.88 |
| 07/12/2018 10:27 | 8.94 |
| 07/12/2018 11:27 | 8.95 |
| 07/12/2018 12:27 | 8.95 |
| 07/12/2018 13:27 | 8.85 |
| 07/12/2018 14:27 | 8.68 |
| 07/12/2018 15:27 | 8.58 |
| 07/12/2018 16:27 | 8.5 |
| 07/12/2018 17:27 | 8.42 |
| 07/12/2018 18:27 | 8.32 |
| 07/12/2018 19:27 | 8.29 |
| 07/12/2018 20:27 | 8.31 |
| 07/12/2018 21:27 | 8.33 |
| 07/12/2018 22:27 | 8.39 |
| 07/12/2018 23:27 | 8.47 |
| 08/12/2018 00:27 | 8.52 |
| 08/12/2018 01:27 | 8.43 |
| 08/12/2018 02:27 | 8.24 |
| 08/12/2018 03:27 | 8.02 |
| 08/12/2018 04:27 | 7.84 |
| 08/12/2018 05:27 | 7.68 |
| 08/12/2018 06:27 | 7.56 |
| 08/12/2018 07:27 | 7.42 |
| 08/12/2018 08:27 | 7.37 |
| 08/12/2018 09:27 | 7.38 |
| 08/12/2018 10:27 | 7.47 |
| 08/12/2018 11:27 | 7.55 |
| 08/12/2018 12:27 | 7.55 |
| 08/12/2018 13:27 | 7.53 |
| 08/12/2018 14:27 | 7.58 |
| 08/12/2018 15:27 | 7.68 |
| 08/12/2018 16:27 | 7.68 |
| 08/12/2018 17:27 | 7.73 |
| 08/12/2018 18:27 | 7.77 |
| 08/12/2018 19:27 | 7.85 |
| 08/12/2018 20:27 | 7.92 |
| 08/12/2018 21:27 | 7.96 |
| 08/12/2018 22:27 | 7.93 |
| 08/12/2018 23:27 | 7.96 |
| 09/12/2018 00:27 | 7.99 |
| 09/12/2018 01:27 | 7.98 |
| 09/12/2018 02:27 | 7.97 |
| 09/12/2018 03:27 | 7.9 |
| 09/12/2018 04:27 | 7.87 |
| 09/12/2018 05:27 | 7.87 |
| 09/12/2018 06:27 | 7.87 |
| 09/12/2018 07:27 | 7.97 |
| 09/12/2018 08:27 | 8.02 |
| 09/12/2018 09:27 | 8.06 |
| 09/12/2018 10:27 | 8.14 |
| 09/12/2018 11:27 | 8.22 |
| 09/12/2018 12:27 | 8.15 |
| 09/12/2018 13:27 | 8.19 |
| 09/12/2018 14:27 | 8.12 |
| 09/12/2018 15:27 | 8.06 |
| 09/12/2018 16:27 | 7.99 |
| 09/12/2018 17:27 | 7.98 |
| 09/12/2018 18:27 | 7.89 |
| 09/12/2018 19:27 | 7.74 |
| 09/12/2018 20:27 | 7.61 |
| 09/12/2018 21:27 | 7.46 |
| 09/12/2018 22:27 | 7.33 |
| 09/12/2018 23:27 | 7.31 |
| 10/12/2018 00:27 | 7.3 |
| 10/12/2018 01:27 | 7.32 |
| 10/12/2018 02:27 | 7.31 |
| 10/12/2018 03:27 | 7.35 |
| 10/12/2018 04:27 | 7.42 |
| 10/12/2018 05:27 | 7.44 |
| 10/12/2018 06:27 | 7.48 |
| 10/12/2018 07:27 | 7.57 |
| 10/12/2018 08:27 | 7.65 |
| 10/12/2018 09:27 | 7.77 |
| 10/12/2018 10:27 | 7.91 |
| 10/12/2018 11:27 | 8.02 |
| 10/12/2018 12:27 | 8.05 |
| 10/12/2018 13:27 | 8.06 |
| 10/12/2018 14:27 | 8.05 |
| 10/12/2018 15:27 | 7.99 |
| 10/12/2018 16:27 | 7.93 |
| 10/12/2018 17:27 | 7.9 |
| 10/12/2018 18:27 | 7.88 |
| 10/12/2018 19:27 | 7.88 |
| 10/12/2018 20:27 | 7.89 |
| 10/12/2018 21:27 | 7.89 |
| 10/12/2018 22:27 | 7.88 |
| 10/12/2018 23:27 | 7.88 |
| 11/12/2018 00:27 | 7.89 |
| 11/12/2018 01:27 | 7.89 |
| 11/12/2018 02:27 | 7.88 |
| 11/12/2018 03:27 | 7.87 |
| 11/12/2018 04:27 | 7.87 |
| 11/12/2018 05:27 | 7.86 |
| 11/12/2018 06:27 | 7.83 |
| 11/12/2018 07:27 | 7.84 |
| 11/12/2018 08:27 | 7.85 |
| 11/12/2018 09:27 | 7.93 |
| 11/12/2018 10:27 | 7.98 |
| 11/12/2018 11:27 | 8.05 |
| 11/12/2018 12:27 | 8.05 |
| 11/12/2018 13:27 | 8.05 |
| 11/12/2018 14:27 | 8.01 |
| 11/12/2018 15:27 | 8.01 |
| 11/12/2018 16:27 | 8.01 |
| 11/12/2018 17:27 | 8 |
| 11/12/2018 18:27 | 8 |
| 11/12/2018 19:27 | 8.05 |
| 11/12/2018 20:27 | 8.09 |
| 11/12/2018 21:27 | 8.14 |
| 11/12/2018 22:27 | 8.19 |
| 11/12/2018 23:27 | 8.26 |
| 12/12/2018 00:27 | 8.31 |
| 12/12/2018 01:27 | 8.37 |
| 12/12/2018 02:27 | 8.43 |
| 12/12/2018 03:27 | 8.51 |
| 12/12/2018 04:27 | 8.55 |
| 12/12/2018 05:27 | 8.6 |
| 12/12/2018 06:27 | 8.6 |
| 12/12/2018 07:27 | 8.58 |
| 12/12/2018 08:27 | 8.62 |
| 12/12/2018 09:27 | 8.74 |
| 12/12/2018 10:27 | 8.9 |
| 12/12/2018 11:27 | 9 |
| 12/12/2018 12:27 | 9.07 |
| 12/12/2018 13:27 | 9.06 |
| 12/12/2018 14:27 | 9.01 |
| 12/12/2018 15:27 | 8.98 |
| 12/12/2018 16:27 | 8.97 |
| 12/12/2018 17:27 | 8.91 |
| 12/12/2018 18:27 | 8.91 |
| 12/12/2018 19:27 | 8.88 |
| 12/12/2018 20:27 | 8.9 |
| 12/12/2018 21:27 | 8.91 |
| 12/12/2018 22:27 | 8.9 |
| 12/12/2018 23:27 | 8.93 |
| 13/12/2018 00:27 | 8.91 |
| 13/12/2018 01:27 | 8.92 |
| 13/12/2018 02:27 | 8.9 |
| 13/12/2018 03:27 | 8.89 |
| 13/12/2018 04:27 | 8.87 |
| 13/12/2018 05:27 | 8.86 |
| 13/12/2018 06:27 | 8.9 |
| 13/12/2018 07:27 | 8.89 |
| 13/12/2018 08:27 | 8.88 |
| 13/12/2018 09:27 | 8.93 |
| 13/12/2018 10:27 | 9.01 |
| 13/12/2018 11:27 | 9.06 |
| 13/12/2018 12:27 | 9.06 |
| 13/12/2018 13:27 | 9.02 |
| 13/12/2018 14:27 | 8.97 |
| 13/12/2018 15:27 | 8.94 |
| 13/12/2018 16:27 | 8.9 |
| 13/12/2018 17:27 | 8.81 |
| 13/12/2018 18:27 | 8.74 |
| 13/12/2018 19:27 | 8.64 |
| 13/12/2018 20:27 | 8.58 |
| 13/12/2018 21:27 | 8.46 |
| 13/12/2018 22:27 | 8.35 |
| 13/12/2018 23:27 | 8.27 |
| 14/12/2018 00:27 | 8.16 |
| 14/12/2018 01:27 | 8.07 |
| 14/12/2018 02:27 | 7.97 |
| 14/12/2018 03:27 | 7.85 |
| 14/12/2018 04:27 | 7.78 |
| 14/12/2018 05:27 | 7.7 |
| 14/12/2018 06:27 | 7.63 |
| 14/12/2018 07:27 | 7.55 |
| 14/12/2018 08:27 | 7.49 |
| 14/12/2018 09:27 | 7.47 |
| 14/12/2018 10:27 | 7.45 |
| 14/12/2018 11:27 | 7.43 |
| 14/12/2018 12:27 | 7.41 |
| 14/12/2018 13:27 | 7.38 |
| 14/12/2018 14:27 | 7.35 |
| 14/12/2018 15:27 | 7.4 |
| 14/12/2018 16:27 | 7.37 |
| 14/12/2018 17:27 | 7.33 |
| 14/12/2018 18:27 | 7.31 |
| 14/12/2018 19:27 | 7.28 |
| 14/12/2018 20:27 | 7.28 |
| 14/12/2018 21:27 | 7.28 |
| 14/12/2018 22:27 | 7.28 |
| 14/12/2018 23:27 | 7.29 |
| 15/12/2018 00:27 | 7.32 |
| 15/12/2018 01:27 | 7.33 |
| 15/12/2018 02:27 | 7.27 |
| 15/12/2018 03:27 | 7.18 |
| 15/12/2018 04:27 | 7.15 |
| 15/12/2018 05:27 | 7.11 |
| 15/12/2018 06:27 | 7.1 |
| 15/12/2018 07:27 | 7.08 |
| 15/12/2018 08:27 | 7.02 |
| 15/12/2018 09:27 | 7.05 |
| 15/12/2018 10:27 | 7.12 |
| 15/12/2018 11:27 | 7.18 |
| 15/12/2018 12:27 | 7.22 |
| 15/12/2018 13:27 | 7.22 |
| 15/12/2018 14:27 | 7.23 |
| 15/12/2018 15:27 | 7.19 |
| 15/12/2018 16:27 | 7.26 |
| 15/12/2018 17:27 | 7.28 |
| 15/12/2018 18:27 | 7.33 |
| 15/12/2018 19:27 | 7.38 |
| 15/12/2018 20:27 | 7.41 |
| 15/12/2018 21:27 | 7.45 |
| 15/12/2018 22:27 | 7.48 |
| 15/12/2018 23:27 | 7.49 |
| 16/12/2018 00:27 | 7.49 |
| 16/12/2018 01:27 | 7.49 |
| 16/12/2018 02:27 | 7.46 |
| 16/12/2018 03:27 | 7.46 |
| 16/12/2018 04:27 | 7.42 |
| 16/12/2018 05:27 | 7.4 |
| 16/12/2018 06:27 | 7.37 |
| 16/12/2018 07:27 | 7.25 |
| 16/12/2018 08:27 | 7.07 |
| 16/12/2018 09:27 | 7.11 |
| 16/12/2018 10:27 | 6.97 |
| 16/12/2018 11:27 | 7.02 |
| 16/12/2018 12:27 | 7.12 |
| 16/12/2018 13:27 | 7.18 |
| 16/12/2018 14:27 | 7.24 |
| 16/12/2018 15:27 | 7.29 |
| 16/12/2018 16:27 | 7.41 |
| 16/12/2018 17:27 | 7.53 |
| 16/12/2018 18:27 | 7.57 |
| 16/12/2018 19:27 | 7.34 |
| 16/12/2018 20:27 | 7.16 |
| 16/12/2018 21:27 | 7.08 |
| 16/12/2018 22:27 | 7.03 |
| 16/12/2018 23:27 | 7.02 |
| 17/12/2018 00:27 | 7.02 |
| 17/12/2018 01:27 | 7.01 |
| 17/12/2018 02:27 | 7 |
| 17/12/2018 03:27 | 7.01 |
| 17/12/2018 04:27 | 6.99 |
| 17/12/2018 05:27 | 6.97 |
| 17/12/2018 06:27 | 6.9 |
| 17/12/2018 07:27 | 6.89 |
| 17/12/2018 08:27 | 6.86 |
| 17/12/2018 09:27 | 6.9 |
| 17/12/2018 10:27 | 6.98 |
| 17/12/2018 11:27 | 7.01 |
| 17/12/2018 12:27 | 7.05 |
| 17/12/2018 13:27 | 7.1 |
| 17/12/2018 14:27 | 7.04 |
| 17/12/2018 15:27 | 7.02 |
| 17/12/2018 16:27 | 6.98 |
| 17/12/2018 17:27 | 6.95 |
| 17/12/2018 18:27 | 6.93 |
| 17/12/2018 19:27 | 6.87 |
| 17/12/2018 20:27 | 6.82 |
| 17/12/2018 21:27 | 6.8 |
| 17/12/2018 22:27 | 6.72 |
| 17/12/2018 23:27 | 6.65 |
| 18/12/2018 00:27 | 6.6 |
| 18/12/2018 01:27 | 6.57 |
| 18/12/2018 02:27 | 6.52 |
| 18/12/2018 03:27 | 6.45 |
| 18/12/2018 04:27 | 6.4 |
| 18/12/2018 05:27 | 6.42 |
| 18/12/2018 06:27 | 6.44 |
| 18/12/2018 07:27 | 6.5 |
| 18/12/2018 08:27 | 6.55 |
| 18/12/2018 09:27 | 6.68 |
| 18/12/2018 10:27 | 6.91 |
| 18/12/2018 11:27 | 7.17 |
| 18/12/2018 12:27 | 7.41 |
| 18/12/2018 13:27 | 7.6 |
| 18/12/2018 14:27 | 7.78 |
| 18/12/2018 15:27 | 7.94 |
| 18/12/2018 16:27 | 8.09 |
| 18/12/2018 17:27 | 8.23 |
| 18/12/2018 18:27 | 8.35 |
| 18/12/2018 19:27 | 8.48 |
| 18/12/2018 20:27 | 8.59 |
| 18/12/2018 21:27 | 8.73 |
| 18/12/2018 22:27 | 8.81 |
| 18/12/2018 23:27 | 8.89 |
| 19/12/2018 00:27 | 8.94 |
| 19/12/2018 01:27 | 8.98 |
| 19/12/2018 02:27 | 9.02 |
| 19/12/2018 03:27 | 9.03 |
| 19/12/2018 04:27 | 9.07 |
| 19/12/2018 05:27 | 8.98 |
| 19/12/2018 06:27 | 8.65 |
| 19/12/2018 07:27 | 8.51 |
| 19/12/2018 08:27 | 8.39 |
| 19/12/2018 09:27 | 8.34 |
| 19/12/2018 10:27 | 8.37 |
| 19/12/2018 11:27 | 8.42 |
| 19/12/2018 12:27 | 8.36 |
| 19/12/2018 13:27 | 8.3 |
| 19/12/2018 14:27 | 8.23 |
| 19/12/2018 15:27 | 8.13 |
| 19/12/2018 16:27 | 8.01 |
| 19/12/2018 17:27 | 7.88 |
| 19/12/2018 18:27 | 7.82 |
| 19/12/2018 19:27 | 7.72 |
| 19/12/2018 20:27 | 7.67 |
| 19/12/2018 21:27 | 7.57 |
| 19/12/2018 22:27 | 7.47 |
| 19/12/2018 23:27 | 7.37 |
| 20/12/2018 00:27 | 7.28 |
| 20/12/2018 01:27 | 7.22 |
| 20/12/2018 02:27 | 7.15 |
| 20/12/2018 03:27 | 7.08 |
| 20/12/2018 04:27 | 7.04 |
| 20/12/2018 05:27 | 7.01 |
| 20/12/2018 06:27 | 7.02 |
| 20/12/2018 07:27 | 7 |
| 20/12/2018 08:27 | 7.03 |
| 20/12/2018 09:27 | 7.18 |
| 20/12/2018 10:27 | 7.32 |
| 20/12/2018 11:27 | 7.41 |
| 20/12/2018 12:27 | 7.5 |
| 20/12/2018 13:27 | 7.58 |
| 20/12/2018 14:27 | 7.59 |
| 20/12/2018 15:27 | 7.63 |
| 20/12/2018 16:27 | 7.67 |
| 20/12/2018 17:27 | 7.63 |
| 20/12/2018 18:27 | 7.55 |
| 20/12/2018 19:27 | 7.54 |
| 20/12/2018 20:27 | 7.59 |
| 20/12/2018 21:27 | 7.61 |
| 20/12/2018 22:27 | 7.65 |
| 20/12/2018 23:27 | 7.67 |
| 21/12/2018 00:27 | 7.68 |
| 21/12/2018 01:27 | 7.65 |
| 21/12/2018 02:27 | 7.65 |
| 21/12/2018 03:27 | 7.6 |
| 21/12/2018 04:27 | 7.61 |
| 21/12/2018 05:27 | 7.64 |
| 21/12/2018 06:27 | 7.63 |
| 21/12/2018 07:27 | 7.62 |
| 21/12/2018 08:27 | 7.67 |
| 21/12/2018 09:27 | 7.73 |
| 21/12/2018 10:27 | 7.85 |
| 21/12/2018 11:27 | 7.97 |
| 21/12/2018 12:27 | 8.03 |
| 21/12/2018 13:27 | 8.05 |
| 21/12/2018 14:27 | 7.98 |
| 21/12/2018 15:27 | 7.89 |
| 21/12/2018 16:27 | 7.87 |
| 21/12/2018 17:27 | 7.86 |
| 21/12/2018 18:27 | 7.85 |
| 21/12/2018 19:27 | 7.79 |
| 21/12/2018 20:27 | 7.72 |
| 21/12/2018 21:27 | 7.69 |
| 21/12/2018 22:27 | 7.68 |
| 21/12/2018 23:27 | 7.68 |
| 22/12/2018 00:27 | 7.65 |
| 22/12/2018 01:27 | 7.62 |
| 22/12/2018 02:27 | 7.63 |
| 22/12/2018 03:27 | 7.63 |
| 22/12/2018 04:27 | 7.65 |
| 22/12/2018 05:27 | 7.64 |
| 22/12/2018 06:27 | 7.68 |
| 22/12/2018 07:27 | 7.74 |
| 22/12/2018 08:27 | 7.78 |
| 22/12/2018 09:27 | 7.89 |
| 22/12/2018 10:27 | 7.96 |
| 22/12/2018 11:27 | 8.12 |
| 22/12/2018 12:27 | 8.15 |
| 22/12/2018 13:27 | 8.12 |
| 22/12/2018 14:27 | 8.05 |
| 22/12/2018 15:27 | 7.98 |
| 22/12/2018 16:27 | 7.92 |
| 22/12/2018 17:27 | 7.89 |
| 22/12/2018 18:27 | 7.87 |
| 22/12/2018 19:27 | 7.86 |
| 22/12/2018 20:27 | 7.84 |
| 22/12/2018 21:27 | 7.84 |
| 22/12/2018 22:27 | 7.76 |
| 22/12/2018 23:27 | 7.73 |
| 23/12/2018 00:27 | 7.72 |
| 23/12/2018 01:27 | 7.75 |
| 23/12/2018 02:27 | 7.77 |
| 23/12/2018 03:27 | 7.74 |
| 23/12/2018 04:27 | 7.68 |
| 23/12/2018 05:27 | 7.65 |
| 23/12/2018 06:27 | 7.61 |
| 23/12/2018 07:27 | 7.6 |
| 23/12/2018 08:27 | 7.6 |
| 23/12/2018 09:27 | 7.63 |
| 23/12/2018 10:27 | 7.68 |
| 23/12/2018 11:27 | 7.78 |
| 23/12/2018 12:27 | 7.85 |
| 23/12/2018 13:27 | 7.83 |
| 23/12/2018 14:27 | 7.75 |
| 23/12/2018 15:27 | 7.63 |
| 23/12/2018 16:27 | 7.57 |
| 23/12/2018 17:27 | 7.46 |
| 23/12/2018 18:27 | 7.35 |
| 23/12/2018 19:27 | 7.33 |
| 23/12/2018 20:27 | 7.32 |
| 23/12/2018 21:27 | 7.32 |
| 23/12/2018 22:27 | 7.42 |
| 23/12/2018 23:27 | 7.46 |
| 24/12/2018 00:27 | 7.48 |
| 24/12/2018 01:27 | 7.48 |
| 24/12/2018 02:27 | 7.47 |
| 24/12/2018 03:27 | 7.45 |
| 24/12/2018 04:27 | 7.41 |
| 24/12/2018 05:27 | 7.44 |
| 24/12/2018 06:27 | 7.46 |
| 24/12/2018 07:27 | 7.48 |
| 24/12/2018 08:27 | 7.48 |
| 24/12/2018 09:27 | 7.53 |
| 24/12/2018 10:27 | 7.55 |
| 24/12/2018 11:27 | 7.52 |
| 24/12/2018 12:27 | 7.54 |
| 24/12/2018 13:27 | 7.56 |
| 24/12/2018 14:27 | 7.55 |
| 24/12/2018 15:27 | 7.45 |
| 24/12/2018 16:27 | 7.29 |
| 24/12/2018 17:27 | 7.12 |
| 24/12/2018 18:27 | 7.1 |
| 24/12/2018 19:27 | 7.05 |
| 24/12/2018 20:27 | 6.93 |
| 24/12/2018 21:27 | 6.81 |
| 24/12/2018 22:27 | 6.64 |
| 24/12/2018 23:27 | 6.63 |
| 25/12/2018 00:27 | 6.6 |
| 25/12/2018 01:27 | 6.55 |
| 25/12/2018 02:27 | 6.41 |
| 25/12/2018 03:27 | 6.23 |
| 25/12/2018 04:27 | 6.09 |
| 25/12/2018 05:27 | 5.95 |
| 25/12/2018 06:27 | 5.83 |
| 25/12/2018 07:27 | 5.69 |
| 25/12/2018 08:27 | 5.64 |
| 25/12/2018 09:27 | 5.71 |
| 25/12/2018 10:27 | 5.9 |
| 25/12/2018 11:27 | 6.04 |
| 25/12/2018 12:27 | 6.14 |
| 25/12/2018 13:27 | 6.19 |
| 25/12/2018 14:27 | 6.21 |
| 25/12/2018 15:27 | 6.33 |
| 25/12/2018 16:27 | 6.45 |
| 25/12/2018 17:27 | 6.57 |
| 25/12/2018 18:27 | 6.71 |
| 25/12/2018 19:27 | 6.9 |
| 25/12/2018 20:27 | 7.05 |
| 25/12/2018 21:27 | 7.22 |
| 25/12/2018 22:27 | 7.38 |
| 25/12/2018 23:27 | 7.5 |
| 26/12/2018 00:27 | 7.59 |
| 26/12/2018 01:27 | 7.72 |
| 26/12/2018 02:27 | 7.78 |
| 26/12/2018 03:27 | 7.87 |
| 26/12/2018 04:27 | 7.93 |
| 26/12/2018 05:27 | 7.99 |
| 26/12/2018 06:27 | 8.08 |
| 26/12/2018 07:27 | 8.14 |
| 26/12/2018 08:27 | 8.27 |
| 26/12/2018 09:27 | 8.43 |
| 26/12/2018 10:27 | 8.53 |
| 26/12/2018 11:27 | 8.63 |
| 26/12/2018 12:27 | 8.76 |
| 26/12/2018 13:27 | 8.82 |
| 26/12/2018 14:27 | 8.8 |
| 26/12/2018 15:27 | 8.72 |
| 26/12/2018 16:27 | 8.66 |
| 26/12/2018 17:27 | 8.62 |
| 26/12/2018 18:27 | 8.59 |
| 26/12/2018 19:27 | 8.57 |
| 26/12/2018 20:27 | 8.53 |
| 26/12/2018 21:27 | 8.49 |
| 26/12/2018 22:27 | 8.48 |
| 26/12/2018 23:27 | 8.46 |
| 27/12/2018 00:27 | 8.45 |
| 27/12/2018 01:27 | 8.45 |
| 27/12/2018 02:27 | 8.41 |
| 27/12/2018 03:27 | 8.43 |
| 27/12/2018 04:27 | 8.44 |
| 27/12/2018 05:27 | 8.42 |
| 27/12/2018 06:27 | 8.36 |
| 27/12/2018 07:27 | 8.37 |
| 27/12/2018 08:27 | 8.42 |
| 27/12/2018 09:27 | 8.47 |
| 27/12/2018 10:27 | 8.56 |
| 27/12/2018 11:27 | 8.64 |
| 27/12/2018 12:27 | 8.68 |
| 27/12/2018 13:27 | 8.71 |
| 27/12/2018 14:27 | 8.65 |
| 27/12/2018 15:27 | 8.66 |
| 27/12/2018 16:27 | 8.61 |
| 27/12/2018 17:27 | 8.59 |
| 27/12/2018 18:27 | 8.55 |
| 27/12/2018 19:27 | 8.5 |
| 27/12/2018 20:27 | 8.46 |
| 27/12/2018 21:27 | 8.41 |
| 27/12/2018 22:27 | 8.28 |
| 27/12/2018 23:27 | 8.17 |
| 28/12/2018 00:27 | 8.05 |
| 28/12/2018 01:27 | 8.01 |
| 28/12/2018 02:27 | 7.99 |
| 28/12/2018 03:27 | 8.03 |
| 28/12/2018 04:27 | 8.12 |
| 28/12/2018 05:27 | 8.14 |
| 28/12/2018 06:27 | 8.21 |
| 28/12/2018 07:27 | 8.24 |
| 28/12/2018 08:27 | 8.35 |
| 28/12/2018 09:27 | 8.45 |
| 28/12/2018 10:27 | 8.57 |
| 28/12/2018 11:27 | 8.65 |
| 28/12/2018 12:27 | 8.7 |
| 28/12/2018 13:27 | 8.72 |
| 28/12/2018 14:27 | 8.73 |
| 28/12/2018 15:27 | 8.7 |
| 28/12/2018 16:27 | 8.68 |
| 28/12/2018 17:27 | 8.66 |
| 28/12/2018 18:27 | 8.66 |
| 28/12/2018 19:27 | 8.62 |
| 28/12/2018 20:27 | 8.59 |
| 28/12/2018 21:27 | 8.58 |
| 28/12/2018 22:27 | 8.59 |
| 28/12/2018 23:27 | 8.52 |
| 29/12/2018 00:27 | 8.48 |
| 29/12/2018 01:27 | 8.49 |
| 29/12/2018 02:27 | 8.47 |
| 29/12/2018 03:27 | 8.52 |
| 29/12/2018 04:27 | 8.53 |
| 29/12/2018 05:27 | 8.62 |
| 29/12/2018 06:27 | 8.65 |
| 29/12/2018 07:27 | 8.62 |
| 29/12/2018 08:27 | 8.57 |
| 29/12/2018 09:27 | 8.47 |
| 29/12/2018 10:27 | 8.46 |
| 29/12/2018 11:27 | 8.44 |
| 29/12/2018 12:27 | 8.37 |
| 29/12/2018 13:27 | 8.3 |
| 29/12/2018 14:27 | 8.17 |
| 29/12/2018 15:27 | 8.02 |
| 29/12/2018 16:27 | 7.87 |
| 29/12/2018 17:27 | 7.84 |
| 29/12/2018 18:27 | 7.92 |
| 29/12/2018 19:27 | 8.03 |
| 29/12/2018 20:27 | 8.19 |
| 29/12/2018 21:27 | 8.38 |
| 29/12/2018 22:27 | 8.62 |
| 29/12/2018 23:27 | 8.81 |
| 30/12/2018 00:27 | 8.97 |
| 30/12/2018 01:27 | 9.05 |
| 30/12/2018 02:27 | 9.04 |
| 30/12/2018 03:27 | 8.97 |
| 30/12/2018 04:27 | 8.9 |
| 30/12/2018 05:27 | 8.81 |
| 30/12/2018 06:27 | 8.66 |
| 30/12/2018 07:27 | 8.61 |
| 30/12/2018 08:27 | 8.59 |
| 30/12/2018 09:27 | 8.67 |
| 30/12/2018 10:27 | 8.76 |
| 30/12/2018 11:27 | 8.79 |
| 30/12/2018 12:27 | 8.85 |
| 30/12/2018 13:27 | 8.87 |
| 30/12/2018 14:27 | 8.83 |
| 30/12/2018 15:27 | 8.73 |
| 30/12/2018 16:27 | 8.63 |
| 30/12/2018 17:27 | 8.57 |
| 30/12/2018 18:27 | 8.52 |
| 30/12/2018 19:27 | 8.5 |
| 30/12/2018 20:27 | 8.49 |
| 30/12/2018 21:27 | 8.49 |
| 30/12/2018 22:27 | 8.55 |
| 30/12/2018 23:27 | 8.71 |
| 31/12/2018 00:27 | 8.85 |
| 31/12/2018 01:27 | 8.97 |
| 31/12/2018 02:27 | 9.07 |
| 31/12/2018 03:27 | 9.05 |
| 31/12/2018 04:27 | 9.06 |
| 31/12/2018 05:27 | 9.04 |
| 31/12/2018 06:27 | 9.03 |
| 31/12/2018 07:27 | 9.02 |
| 31/12/2018 08:27 | 9.07 |
| 31/12/2018 09:27 | 9.15 |
| 31/12/2018 10:27 | 9.26 |
| 31/12/2018 11:27 | 9.34 |
| 31/12/2018 12:27 | 9.35 |
| 31/12/2018 13:27 | 9.35 |
| 31/12/2018 14:27 | 9.26 |
| 31/12/2018 15:27 | 9.22 |
| 31/12/2018 16:27 | 9.2 |
| 31/12/2018 17:27 | 9.16 |
| 31/12/2018 18:27 | 9.12 |
| 31/12/2018 19:27 | 9.11 |
| 31/12/2018 20:27 | 9.07 |
| 31/12/2018 21:27 | 9.07 |
| 31/12/2018 22:27 | 9.01 |
| 31/12/2018 23:27 | 9.01 |
| 01/01/2019 00:27 | 8.98 |
| 01/01/2019 01:27 | 8.95 |
| 01/01/2019 02:27 | 8.93 |
| 01/01/2019 03:27 | 8.92 |
| 01/01/2019 04:27 | 8.89 |
| 01/01/2019 05:27 | 8.89 |
| 01/01/2019 06:27 | 8.87 |
| 01/01/2019 07:27 | 8.88 |
| 01/01/2019 08:27 | 8.89 |
| 01/01/2019 09:27 | 8.89 |
| 01/01/2019 10:27 | 8.9 |
| 01/01/2019 11:27 | 8.88 |
| 01/01/2019 12:27 | 8.85 |
| 01/01/2019 13:27 | 8.82 |
| 01/01/2019 14:27 | 8.81 |
| 01/01/2019 15:27 | 8.81 |
| 01/01/2019 16:27 | 8.86 |
| 01/01/2019 17:27 | 8.87 |
| 01/01/2019 18:27 | 8.9 |
| 01/01/2019 19:27 | 8.94 |
| 01/01/2019 20:27 | 8.96 |
| 01/01/2019 21:27 | 9 |
| 01/01/2019 22:27 | 9.06 |
| 01/01/2019 23:27 | 9.11 |
| 02/01/2019 00:27 | 9.16 |
| 02/01/2019 01:27 | 9.1 |
| 02/01/2019 02:27 | 9.03 |
| 02/01/2019 03:27 | 8.91 |
| 02/01/2019 04:27 | 8.76 |
| 02/01/2019 05:27 | 8.56 |
| 02/01/2019 06:27 | 8.34 |
| 02/01/2019 07:27 | 8.2 |
| 02/01/2019 08:27 | 8.12 |
| 02/01/2019 09:27 | 8.18 |
| 02/01/2019 10:27 | 8.23 |
| 02/01/2019 11:27 | 8.29 |
| 02/01/2019 12:27 | 8.33 |
| 02/01/2019 13:27 | 8.28 |
| 02/01/2019 14:27 | 8.26 |
| 02/01/2019 15:27 | 8.2 |
| 02/01/2019 16:27 | 8.12 |
| 02/01/2019 17:27 | 8 |
| 02/01/2019 18:27 | 7.78 |
| 02/01/2019 19:27 | 7.5 |
| 02/01/2019 20:27 | 7.14 |
| 02/01/2019 21:27 | 6.8 |
| 02/01/2019 22:27 | 6.57 |
| 02/01/2019 23:27 | 6.48 |
| 03/01/2019 00:27 | 6.44 |
| 03/01/2019 01:27 | 6.45 |
| 03/01/2019 02:27 | 6.49 |
| 03/01/2019 03:27 | 6.56 |
| 03/01/2019 04:27 | 6.64 |
| 03/01/2019 05:27 | 6.6 |
| 03/01/2019 06:27 | 6.52 |
| 03/01/2019 07:27 | 6.51 |
| 03/01/2019 08:27 | 6.51 |
| 03/01/2019 09:27 | 6.54 |
| 03/01/2019 10:27 | 6.61 |
| 03/01/2019 11:27 | 6.73 |
| 03/01/2019 12:27 | 6.79 |
| 03/01/2019 13:27 | 6.72 |
| 03/01/2019 14:27 | 6.61 |
| 03/01/2019 15:27 | 6.43 |
| 03/01/2019 16:27 | 6.27 |
| 03/01/2019 17:27 | 6.17 |
| 03/01/2019 18:27 | 6.21 |
| 03/01/2019 19:27 | 6.23 |
| 03/01/2019 20:27 | 6.3 |
| 03/01/2019 21:27 | 6.38 |
| 03/01/2019 22:27 | 6.48 |
| 03/01/2019 23:27 | 6.58 |
| 04/01/2019 00:27 | 6.65 |
| 04/01/2019 01:27 | 6.73 |
| 04/01/2019 02:27 | 6.83 |
| 04/01/2019 03:27 | 6.87 |
| 04/01/2019 04:27 | 6.94 |
| 04/01/2019 05:27 | 6.97 |
| 04/01/2019 06:27 | 7.01 |
| 04/01/2019 07:27 | 7.08 |
| 04/01/2019 08:27 | 7.14 |
| 04/01/2019 09:27 | 7.24 |
| 04/01/2019 10:27 | 7.35 |
| 04/01/2019 11:27 | 7.45 |
| 04/01/2019 12:27 | 7.55 |
| 04/01/2019 13:27 | 7.6 |
| 04/01/2019 14:27 | 7.6 |
| 04/01/2019 15:27 | 7.58 |
| 04/01/2019 16:27 | 7.55 |
| 04/01/2019 17:27 | 7.55 |
| 04/01/2019 18:27 | 7.56 |
| 04/01/2019 19:27 | 7.56 |
| 04/01/2019 20:27 | 7.56 |
| 04/01/2019 21:27 | 7.56 |
| 04/01/2019 22:27 | 7.56 |
| 04/01/2019 23:27 | 7.58 |
| 05/01/2019 00:27 | 7.61 |
| 05/01/2019 01:27 | 7.6 |
| 05/01/2019 02:27 | 7.61 |
| 05/01/2019 03:27 | 7.59 |
| 05/01/2019 04:27 | 7.57 |
| 05/01/2019 05:27 | 7.56 |
| 05/01/2019 06:27 | 7.55 |
| 05/01/2019 07:27 | 7.55 |
| 05/01/2019 08:27 | 7.54 |
| 05/01/2019 09:27 | 7.58 |
| 05/01/2019 10:27 | 7.58 |
| 05/01/2019 11:27 | 7.59 |
| 05/01/2019 12:27 | 7.57 |
| 05/01/2019 13:27 | 7.54 |
| 05/01/2019 14:27 | 7.47 |
| 05/01/2019 15:27 | 7.43 |
| 05/01/2019 16:27 | 7.35 |
| 05/01/2019 17:27 | 7.3 |
| 05/01/2019 18:27 | 7.24 |
| 05/01/2019 19:27 | 7.17 |
| 05/01/2019 20:27 | 7.11 |
| 05/01/2019 21:27 | 7.07 |
| 05/01/2019 22:27 | 7.03 |
| 05/01/2019 23:27 | 7.02 |
| 06/01/2019 00:27 | 7 |
| 06/01/2019 01:27 | 6.95 |
| 06/01/2019 02:27 | 6.81 |
| 06/01/2019 03:27 | 6.59 |
| 06/01/2019 04:27 | 6.31 |
| 06/01/2019 05:27 | 6.14 |
| 06/01/2019 06:27 | 6.08 |
| 06/01/2019 07:27 | 6.1 |
| 06/01/2019 08:27 | 6.23 |
| 06/01/2019 09:27 | 6.41 |
| 06/01/2019 10:27 | 6.65 |
| 06/01/2019 11:27 | 6.86 |
| 06/01/2019 12:27 | 7.01 |
| 06/01/2019 13:27 | 7.08 |
| 06/01/2019 14:27 | 7.09 |
| 06/01/2019 15:27 | 7.11 |
| 06/01/2019 16:27 | 7.15 |
| 06/01/2019 17:27 | 7.16 |
| 06/01/2019 18:27 | 7.15 |
| 06/01/2019 19:27 | 7.22 |
| 06/01/2019 20:27 | 7.25 |
| 06/01/2019 21:27 | 7.28 |
| 06/01/2019 22:27 | 7.33 |
| 06/01/2019 23:27 | 7.41 |
| 07/01/2019 00:27 | 7.48 |
| 07/01/2019 01:27 | 7.54 |
| 07/01/2019 02:27 | 7.61 |
| 07/01/2019 03:27 | 7.67 |
| 07/01/2019 04:27 | 7.73 |
| 07/01/2019 05:27 | 7.78 |
| 07/01/2019 06:27 | 7.82 |
| 07/01/2019 07:27 | 7.92 |
| 07/01/2019 08:27 | 7.98 |
| 07/01/2019 09:27 | 8.11 |
| 07/01/2019 10:27 | 8.21 |
| 07/01/2019 11:27 | 8.32 |
| 07/01/2019 12:27 | 8.43 |
| 07/01/2019 13:27 | 8.5 |
| 07/01/2019 14:27 | 8.52 |
| 07/01/2019 15:27 | 8.56 |
| 07/01/2019 16:27 | 8.53 |
| 07/01/2019 17:27 | 8.53 |
| 07/01/2019 18:27 | 8.55 |
| 07/01/2019 19:27 | 8.56 |
| 07/01/2019 20:27 | 8.59 |
| 07/01/2019 21:27 | 8.65 |
| 07/01/2019 22:27 | 8.71 |
| 07/01/2019 23:27 | 8.76 |
| 08/01/2019 00:27 | 8.81 |
| 08/01/2019 01:27 | 8.9 |
| 08/01/2019 02:27 | 9 |
| 08/01/2019 03:27 | 9.1 |
| 08/01/2019 04:27 | 9.14 |
| 08/01/2019 05:27 | 9.15 |
| 08/01/2019 06:27 | 9.19 |
| 08/01/2019 07:27 | 9.21 |
| 08/01/2019 08:27 | 9.22 |
| 08/01/2019 09:27 | 9.12 |
| 08/01/2019 10:27 | 9.04 |
| 08/01/2019 11:27 | 8.95 |
| 08/01/2019 12:27 | 8.82 |
| 08/01/2019 13:27 | 8.62 |
| 08/01/2019 14:27 | 8.35 |
| 08/01/2019 15:27 | 8.13 |
| 08/01/2019 16:27 | 7.93 |
| 08/01/2019 17:27 | 7.74 |
| 08/01/2019 18:27 | 7.63 |
| 08/01/2019 19:27 | 7.56 |
| 08/01/2019 20:27 | 7.53 |
| 08/01/2019 21:27 | 7.48 |
| 08/01/2019 22:27 | 7.48 |
| 08/01/2019 23:27 | 7.48 |
| 09/01/2019 00:27 | 7.41 |
| 09/01/2019 01:27 | 7.28 |
| 09/01/2019 02:27 | 7.2 |
| 09/01/2019 03:27 | 7.11 |
| 09/01/2019 04:27 | 7.04 |
| 09/01/2019 05:27 | 7.01 |
| 09/01/2019 06:27 | 7.02 |
| 09/01/2019 07:27 | 7.04 |
| 09/01/2019 08:27 | 7.09 |
| 09/01/2019 09:27 | 7.17 |
| 09/01/2019 10:27 | 7.37 |
| 09/01/2019 11:27 | 7.52 |
| 09/01/2019 12:27 | 7.57 |
| 09/01/2019 13:27 | 7.52 |
| 09/01/2019 14:27 | 7.46 |
| 09/01/2019 15:27 | 7.44 |
| 09/01/2019 16:27 | 7.39 |
| 09/01/2019 17:27 | 7.38 |
| 09/01/2019 18:27 | 7.28 |
| 09/01/2019 19:27 | 7.19 |
| 09/01/2019 20:27 | 7.12 |
| 09/01/2019 21:27 | 7.14 |
| 09/01/2019 22:27 | 7.08 |
| 09/01/2019 23:27 | 7.01 |
| 10/01/2019 00:27 | 6.88 |
| 10/01/2019 01:27 | 6.79 |
| 10/01/2019 02:27 | 6.68 |
| 10/01/2019 03:27 | 6.54 |
| 10/01/2019 04:27 | 6.5 |
| 10/01/2019 05:27 | 6.42 |
| 10/01/2019 06:27 | 6.31 |
| 10/01/2019 07:27 | 6.35 |
| 10/01/2019 08:27 | 6.4 |
| 10/01/2019 09:27 | 6.59 |
| 10/01/2019 10:27 | 6.84 |
| 10/01/2019 11:27 | 7.09 |
| 10/01/2019 12:27 | 7.32 |
| 10/01/2019 13:27 | 7.45 |
| 10/01/2019 14:27 | 7.59 |
| 10/01/2019 15:27 | 7.64 |
| 10/01/2019 16:27 | 7.68 |
| 10/01/2019 17:27 | 7.72 |
| 10/01/2019 18:27 | 7.78 |
| 10/01/2019 19:27 | 7.77 |
| 10/01/2019 20:27 | 7.81 |
| 10/01/2019 21:27 | 7.83 |
| 10/01/2019 22:27 | 7.85 |
| 10/01/2019 23:27 | 7.87 |
| 11/01/2019 00:27 | 7.98 |
| 11/01/2019 01:27 | 8.14 |
| 11/01/2019 02:27 | 8.28 |
| 11/01/2019 03:27 | 8.35 |
| 11/01/2019 04:27 | 8.38 |
| 11/01/2019 05:27 | 8.36 |
| 11/01/2019 06:27 | 8.35 |
| 11/01/2019 07:27 | 8.31 |
| 11/01/2019 08:27 | 8.34 |
| 11/01/2019 09:27 | 8.41 |
| 11/01/2019 10:27 | 8.5 |
| 11/01/2019 11:27 | 8.58 |
| 11/01/2019 12:27 | 8.59 |
| 11/01/2019 13:27 | 8.59 |
| 11/01/2019 14:27 | 8.57 |
| 11/01/2019 15:27 | 8.5 |
| 11/01/2019 16:27 | 8.43 |
| 11/01/2019 17:27 | 8.37 |
| 11/01/2019 18:27 | 8.29 |
| 11/01/2019 19:27 | 8.24 |
| 11/01/2019 20:27 | 8.18 |
| 11/01/2019 21:27 | 8.16 |
| 11/01/2019 22:27 | 8.15 |
| 11/01/2019 23:27 | 8.12 |
| 12/01/2019 00:27 | 8.14 |
| 12/01/2019 01:27 | 8.13 |
| 12/01/2019 02:27 | 8.14 |
| 12/01/2019 03:27 | 8.12 |
| 12/01/2019 04:27 | 8.12 |
| 12/01/2019 05:27 | 8.09 |
| 12/01/2019 06:27 | 8.09 |
| 12/01/2019 07:27 | 8.12 |
| 12/01/2019 08:27 | 8.19 |
| 12/01/2019 09:27 | 8.31 |
| 12/01/2019 10:27 | 8.48 |
| 12/01/2019 11:27 | 8.62 |
| 12/01/2019 12:27 | 8.67 |
| 12/01/2019 13:27 | 8.73 |
| 12/01/2019 14:27 | 8.66 |
| 12/01/2019 15:27 | 8.58 |
| 12/01/2019 16:27 | 8.48 |
| 12/01/2019 17:27 | 8.41 |
| 12/01/2019 18:27 | 8.42 |
| 12/01/2019 19:27 | 8.37 |
| 12/01/2019 20:27 | 8.41 |
| 12/01/2019 21:27 | 8.46 |
| 12/01/2019 22:27 | 8.45 |
| 12/01/2019 23:27 | 8.44 |
| 13/01/2019 00:27 | 8.35 |
| 13/01/2019 01:27 | 8.29 |
| 13/01/2019 02:27 | 8.26 |
| 13/01/2019 03:27 | 8.15 |
| 13/01/2019 04:27 | 8.06 |
| 13/01/2019 05:27 | 8 |
| 13/01/2019 06:27 | 7.94 |
| 13/01/2019 07:27 | 7.91 |
| 13/01/2019 08:27 | 7.99 |
| 13/01/2019 09:27 | 8.14 |
| 13/01/2019 10:27 | 8.42 |
| 13/01/2019 11:27 | 8.8 |
| 13/01/2019 12:27 | 9.15 |
| 13/01/2019 13:27 | 9.22 |
| 13/01/2019 14:27 | 9.2 |
| 13/01/2019 15:27 | 9.06 |
| 13/01/2019 16:27 | 8.96 |
| 13/01/2019 17:27 | 8.87 |
| 13/01/2019 18:27 | 8.77 |
| 13/01/2019 19:27 | 8.7 |
| 13/01/2019 20:27 | 8.6 |
| 13/01/2019 21:27 | 8.58 |
| 13/01/2019 22:27 | 8.6 |
| 13/01/2019 23:27 | 8.62 |
| 14/01/2019 00:27 | 8.65 |
| 14/01/2019 01:27 | 8.7 |
| 14/01/2019 02:27 | 8.72 |
| 14/01/2019 03:27 | 8.79 |
| 14/01/2019 04:27 | 8.86 |
| 14/01/2019 05:27 | 8.92 |
| 14/01/2019 06:27 | 8.98 |
| 14/01/2019 07:27 | 8.98 |
| 14/01/2019 08:27 | 8.98 |
| 14/01/2019 09:27 | 9.05 |
| 14/01/2019 10:27 | 9.01 |
| 14/01/2019 11:27 | 8.97 |
| 14/01/2019 12:27 | 8.87 |
| 14/01/2019 13:27 | 8.73 |
| 14/01/2019 14:27 | 8.51 |
| 14/01/2019 15:27 | 8.32 |
| 14/01/2019 16:27 | 8.13 |
| 14/01/2019 17:27 | 7.94 |
| 14/01/2019 18:27 | 7.82 |
| 14/01/2019 19:27 | 7.71 |
| 14/01/2019 20:27 | 7.67 |
| 14/01/2019 21:27 | 7.64 |
| 14/01/2019 22:27 | 7.53 |
| 14/01/2019 23:27 | 7.45 |
| 15/01/2019 00:27 | 7.28 |
| 15/01/2019 01:27 | 7.17 |
| 15/01/2019 02:27 | 7.1 |
| 15/01/2019 03:27 | 7.11 |
| 15/01/2019 04:27 | 7.14 |
| 15/01/2019 05:27 | 7.22 |
| 15/01/2019 06:27 | 7.28 |
| 15/01/2019 07:27 | 7.31 |
| 15/01/2019 08:27 | 7.4 |
| 15/01/2019 09:27 | 7.59 |
| 15/01/2019 10:27 | 7.77 |
| 15/01/2019 11:27 | 7.97 |
| 15/01/2019 12:27 | 8.07 |
| 15/01/2019 13:27 | 8.14 |
| 15/01/2019 14:27 | 8.16 |
| 15/01/2019 15:27 | 8.12 |
| 15/01/2019 16:27 | 8.12 |
| 15/01/2019 17:27 | 8.12 |
| 15/01/2019 18:27 | 8.08 |
| 15/01/2019 19:27 | 8.02 |
| 15/01/2019 20:27 | 7.99 |
| 15/01/2019 21:27 | 7.97 |
| 15/01/2019 22:27 | 7.94 |
| 15/01/2019 23:27 | 7.99 |
| 16/01/2019 00:27 | 8.01 |
| 16/01/2019 01:27 | 8.03 |
| 16/01/2019 02:27 | 8.03 |
| 16/01/2019 03:27 | 8.05 |
| 16/01/2019 04:27 | 8.04 |
| 16/01/2019 05:27 | 7.96 |
| 16/01/2019 06:27 | 7.93 |
| 16/01/2019 07:27 | 7.93 |
| 16/01/2019 08:27 | 8.02 |
| 16/01/2019 09:27 | 8.14 |
| 16/01/2019 10:27 | 8.27 |
| 16/01/2019 11:27 | 8.42 |
| 16/01/2019 12:27 | 8.51 |
| 16/01/2019 13:27 | 8.57 |
| 16/01/2019 14:27 | 8.6 |
| 16/01/2019 15:27 | 8.59 |
| 16/01/2019 16:27 | 8.57 |
| 16/01/2019 17:27 | 8.55 |
| 16/01/2019 18:27 | 8.49 |
| 16/01/2019 19:27 | 8.43 |
| 16/01/2019 20:27 | 8.36 |
| 16/01/2019 21:27 | 8.28 |
| 16/01/2019 22:27 | 8.22 |
| 16/01/2019 23:27 | 8.19 |
| 17/01/2019 00:27 | 8.14 |
| 17/01/2019 01:27 | 8.05 |
| 17/01/2019 02:27 | 7.97 |
| 17/01/2019 03:27 | 7.82 |
| 17/01/2019 04:27 | 7.61 |
| 17/01/2019 05:27 | 7.35 |
| 17/01/2019 06:27 | 7.12 |
| 17/01/2019 07:27 | 6.88 |
| 17/01/2019 08:27 | 6.83 |
| 17/01/2019 09:27 | 6.89 |
| 17/01/2019 10:27 | 7.12 |
| 17/01/2019 11:27 | 7.08 |
| 17/01/2019 12:27 | 7.03 |
| 17/01/2019 13:27 | 6.88 |
| 17/01/2019 14:27 | 6.87 |
| 17/01/2019 15:27 | 6.74 |
| 17/01/2019 16:27 | 6.54 |
| 17/01/2019 17:27 | 6.57 |
| 17/01/2019 18:27 | 5.94 |
| 17/01/2019 19:27 | 5.16 |
| 17/01/2019 20:27 | 5.46 |
| 17/01/2019 21:27 | 6.04 |
| 17/01/2019 22:27 | 6.08 |
| 17/01/2019 23:27 | 5.94 |
| 18/01/2019 00:27 | 5.89 |
| 18/01/2019 01:27 | 5.87 |
| 18/01/2019 02:27 | 5.81 |
| 18/01/2019 03:27 | 5.72 |
| 18/01/2019 04:27 | 5.68 |
| 18/01/2019 05:27 | 5.69 |
| 18/01/2019 06:27 | 5.68 |
| 18/01/2019 07:27 | 5.7 |
| 18/01/2019 08:27 | 5.8 |
| 18/01/2019 09:27 | 6.07 |
| 18/01/2019 10:27 | 6.28 |
| 18/01/2019 11:27 | 6.43 |
| 18/01/2019 12:27 | 6.56 |
| 18/01/2019 13:27 | 6.63 |
| 18/01/2019 14:27 | 6.58 |
| 18/01/2019 15:27 | 6.46 |
| 18/01/2019 16:27 | 6.3 |
| 18/01/2019 17:27 | 6.16 |
| 18/01/2019 18:27 | 6 |
| 18/01/2019 19:27 | 5.87 |
| 18/01/2019 20:27 | 5.72 |
| 18/01/2019 21:27 | 5.61 |
| 18/01/2019 22:27 | 5.53 |
| 18/01/2019 23:27 | 5.48 |
| 19/01/2019 00:27 | 5.42 |
| 19/01/2019 01:27 | 5.37 |
| 19/01/2019 02:27 | 5.39 |
| 19/01/2019 03:27 | 5.42 |
| 19/01/2019 04:27 | 5.49 |
| 19/01/2019 05:27 | 5.59 |
| 19/01/2019 06:27 | 5.75 |
| 19/01/2019 07:27 | 5.92 |
| 19/01/2019 08:27 | 6.13 |
| 19/01/2019 09:27 | 6.28 |
| 19/01/2019 10:27 | 6.4 |
| 19/01/2019 11:27 | 6.52 |
| 19/01/2019 12:27 | 6.66 |
| 19/01/2019 13:27 | 6.74 |
| 19/01/2019 14:27 | 6.8 |
| 19/01/2019 15:27 | 6.86 |
| 19/01/2019 16:27 | 6.91 |
| 19/01/2019 17:27 | 6.95 |
| 19/01/2019 18:27 | 7 |
| 19/01/2019 19:27 | 7.01 |
| 19/01/2019 20:27 | 7 |
| 19/01/2019 21:27 | 6.91 |
| 19/01/2019 22:27 | 6.8 |
| 19/01/2019 23:27 | 6.7 |
| 20/01/2019 00:27 | 6.67 |
| 20/01/2019 01:27 | 6.73 |
| 20/01/2019 02:27 | 6.73 |
| 20/01/2019 03:27 | 6.66 |
| 20/01/2019 04:27 | 6.56 |
| 20/01/2019 05:27 | 6.55 |
| 20/01/2019 06:27 | 6.58 |
| 20/01/2019 07:27 | 6.66 |
| 20/01/2019 08:27 | 6.83 |
| 20/01/2019 09:27 | 7 |
| 20/01/2019 10:27 | 7.17 |
| 20/01/2019 11:27 | 7.31 |
| 20/01/2019 12:27 | 7.41 |
| 20/01/2019 13:27 | 7.48 |
| 20/01/2019 14:27 | 7.47 |
| 20/01/2019 15:27 | 7.41 |
| 20/01/2019 16:27 | 7.26 |
| 20/01/2019 17:27 | 7.11 |
| 20/01/2019 18:27 | 6.98 |
| 20/01/2019 19:27 | 6.81 |
| 20/01/2019 20:27 | 6.82 |
| 20/01/2019 21:27 | 6.76 |
| 20/01/2019 22:27 | 6.82 |
| 20/01/2019 23:27 | 6.81 |
| 21/01/2019 00:27 | 6.86 |
| 21/01/2019 01:27 | 6.91 |
| 21/01/2019 02:27 | 6.9 |
| 21/01/2019 03:27 | 6.88 |
| 21/01/2019 04:27 | 6.86 |
| 21/01/2019 05:27 | 6.85 |
| 21/01/2019 06:27 | 6.94 |
| 21/01/2019 07:27 | 7.01 |
| 21/01/2019 08:27 | 6.99 |
| 21/01/2019 09:27 | 7.05 |
| 21/01/2019 10:27 | 7.18 |
| 21/01/2019 11:27 | 7.25 |
| 21/01/2019 12:27 | 7.28 |
| 21/01/2019 13:27 | 7.13 |
| 21/01/2019 14:27 | 7.02 |
| 21/01/2019 15:27 | 6.83 |
| 21/01/2019 16:27 | 6.65 |
| 21/01/2019 17:27 | 6.54 |
| 21/01/2019 18:27 | 6.46 |
| 21/01/2019 19:27 | 6.31 |
| 21/01/2019 20:27 | 6.11 |
| 21/01/2019 21:27 | 5.93 |
| 21/01/2019 22:27 | 5.75 |
| 21/01/2019 23:27 | 5.54 |
| 22/01/2019 00:27 | 5.41 |
| 22/01/2019 01:27 | 5.3 |
| 22/01/2019 02:27 | 5.28 |
| 22/01/2019 03:27 | 5.28 |
| 22/01/2019 04:27 | 5.31 |
| 22/01/2019 05:27 | 5.31 |
| 22/01/2019 06:27 | 5.33 |
| 22/01/2019 07:27 | 5.34 |
| 22/01/2019 08:27 | 5.4 |
| Average Temp | 7.721849 |
| Standard Deviation | 0.845312 |

Supplementary Table 2: Biofilm biomass (Ash Free Dry Weight), Respiration, Net Primary Production and Gross Primary Production data for each individual Contaminant Exposure Experiment.

| Treatment | Ash Free Dry Weight mg cm^-2^ | Respiration μg cm^-2^ h^-1^ | NPP μg cm^-2^ h^-1^ | GPP μg cm^-2^ h^-1^ |
| --- | --- | --- | --- | --- |
| Ibuprofen | 2.012412497 | -67.25097037 | -46.91825772 | 20.33271266 |
| Ibuprofen | 3.370790933 | -44.94536995 | -90.58670753 | -45.64133758 |
| Ibuprofen | 4.993298509 | -68.52779542 | -75.03379069 | -6.505995265 |
| Ibuprofen | 16.31311881 | -67.9548575 | -55.341295 | 12.6135625 |
| Ibuprofen | 3.314191832 | -48.97790404 | -75.24933166 | -26.27142762 |
| Ibuprofen | 1.93065824 | -49.19782085 | -24.13298113 | 25.06483972 |
| Ibuprofen | 1.452710272 | -68.51313877 | -122.6843763 | -54.17123757 |
| Ibuprofen | 1.484154217 | -56.04029114 | -82.29499293 | -26.25470179 |
| Ibuprofen | 1.257757811 | -52.93531702 | -58.58695392 | -5.651636893 |
| Ibuprofen | 1.113115663 | -67.80286174 | -12.64952177 | 55.15333997 |
| EE2 | 3.433678824 | -107.121795 | -114.7034295 | -7.581634511 |
| EE2 | 2.490360466 | -87.58486886 | -86.30982295 | 1.275045907 |
| EE2 | 9.502360261 | -114.0893949 | -102.402228 | 11.68716685 |
| EE2 | 1.446421483 | -108.4957225 | -88.26257129 | 20.23315117 |
| EE2 | 2.012412497 | -84.42767141 | -90.52935065 | -6.101679243 |
| EE2 | 0.987339882 | -117.7424407 | -61.59616753 | 56.1462732 |
| EE2 | 2.150765857 | -118.3025067 | -105.1427994 | 13.15970731 |
| EE2 | 3.068929059 | -109.0157673 | -118.9656928 | -9.949925579 |
| EE2 | 9.408028425 | -115.3171161 | -127.7452869 | -12.42817078 |
| EE2 | 1.591063631 | -110.1432762 | -86.40400301 | 23.73927319 |
| Ibuprofen + EE2 | 1.100538085 | -85.2170475 | -25.64013186 | 59.57691564 |
| Ibuprofen + EE2 | 1.540753318 | -92.74568805 | -105.773808 | -13.02811999 |
| Ibuprofen + EE2 | 1.408688748 | -113.0905582 | -94.6162405 | 18.47431766 |
| Ibuprofen + EE2 | 0.886719257 | -115.1686542 | -101.1195541 | 14.0491001 |
| Ibuprofen + EE2 | 1.999834919 | -85.93866336 | -26.38191104 | 59.55675232 |
| Ibuprofen + EE2 | 4.72288058 | -97.002267 | -9.229376766 | 87.77289023 |
| Ibuprofen + EE2 | 5.301449173 | -90.31942755 | -141.2279123 | -50.90848473 |
| Ibuprofen + EE2 | 2.635002614 | -105.9254687 | -128.998114 | -23.07264525 |
| Ibuprofen + EE2 | 1.66024031 | -84.6005067 | -58.02803982 | 26.57246688 |
| Ibuprofen + EE2 | 2.361789667 | -119.4346833 | -35.9011583 | 83.533525 |
| Control | 4.012247417 | -85.78892393 | -87.35580644 | -1.566882511 |
| Control | 3.219859996 | -102.9915941 | -123.0940459 | -20.1024518 |
| Control | 9.684735144 | -93.33146213 | -59.93219048 | 33.39927166 |
| Control | 0.874141679 | -104.9807215 | -75.79220999 | 29.18851148 |
| Control | 1.069094139 | -96.99917247 | -81.92655996 | 15.07261251 |
| Control | 0.389904921 | -97.43013533 | -100.1487064 | -2.718571113 |
| Control | 1.352089647 | -138.6673007 | -91.82145936 | 46.84584136 |
| Control | 1.364667225 | -93.76797192 | -108.7048853 | -14.93691335 |
| Control | 3.710385542 | -93.76419663 | -117.0503767 | -23.28618006 |
| Control | 2.490360466 | -110.8301289 | -91.90078345 | 18.92934546 |
